# A histone acetylome-wide association study of Alzheimer’s disease: neuropathology-associated regulatory variation in the human entorhinal cortex

**DOI:** 10.1101/183541

**Authors:** Sarah J. Marzi, Teodora Ribarska, Adam R. Smith, Eilis Hannon, Jeremie Poschmann, Karen Moore, Claire Troakes, Safa Al-Sarraj, Stephan Beck, Stuart Newman, Katie Lunnon, Leonard C. Schalkwyk, Jonathan Mill

## Abstract

Alzheimer’s disease (AD) is a chronic neurodegenerative disorder characterized by the progressive accumulation of amyloid-β (Aβ) plaques and neurofibrillary tangles in the neocortex. Recent studies have implicated a role for regulatory genomic variation in AD progression, finding widespread evidence for altered DNA methylation associated with neuropathology. To date, however, no study has systematically examined other types of regulatory genomic modifications in AD. In this study, we quantified genome-wide patterns of lysine H3K27 acetylation (H3K27ac) - a robust mark of active enhancers and promoters that is strongly correlated with gene expression and transcription factor binding - in entorhinal cortex samples from AD cases and matched controls (n = 47) using chromatin immunoprecipitation followed by highly parallel sequencing (ChIP-seq). Across ~182,000 robustly detected H3K27ac peak regions, we found widespread acetylomic variation associated with AD neuropathology, identifying 4,162 differential peaks (FDR < 0.05) between AD cases and controls. These differentially acetylated peaks are enriched in disease-specific biological pathways and include regions annotated to multiple genes directly involved in the progression of Aβ and tau pathology (e.g. *APP*, *PSEN1*, *PSEN2*, *MAPT*), as well as genomic regions containing variants associated with sporadic late-onset AD. This is the first study of variable H3K27ac yet undertaken in AD and the largest study investigating this modification in the entorhinal cortex. In addition to identifying molecular pathways associated with AD neuropathology, we present a framework for genome-wide studies of histone modifications in complex disease, integrating our data with results obtained from genome-wide association studies as well as other epigenetic marks profiled on the same samples.

---

Alzheimer’s disease (AD) is a chronic neurodegenerative disorder characterized by cognitive decline and memory loss that contributes substantially to the global burden of disease, affecting in excess of 26 million people worldwide^1^. The symptoms of AD are associated with progressive neuropathology in the neocortex, with regions surrounding the entorhinal cortex being particularly affected early in the disease^2^. These neuropathological hallmarks of AD include the extracellular deposition of neurotoxic amyloid-β (Aβ) in the form of amyloid plaques and an accumulation of intracellular neurofibrillary tangles composed of hyperphosphorylated tau^3^. Despite progress in understanding risk factors contributing to AD progression, the mechanisms involved in disease progression are not fully understood and long-term treatments, reversing the cellular disease process in the cortex, are elusive.

There has been considerable success in identifying genetic risk factors for AD^4^. While autosomal dominant mutations in three genes (*APP*, *PSEN1*, and *PSEN2*) can explain early-onset (< 65 years) familial AD, these account for only 1-5% of the total disease burden^5^. Most cases of AD are late-onset (> 65 years), non-Mendelian and highly sporadic, with susceptibility attributed to the action of highly prevalent genetic variants of low penetrance. In addition to the well-established risk associated with the *APOE* locus^6, 7^ there has been notable success in identifying novel AD-associated variants capitalising on the power of genome-wide association studies (GWAS) in large sample cohorts; a recent large GWAS meta-analysis of AD, incorporating > 74,000 samples, identified 19 genome-wide significant risk loci for sporadic AD^8^. Despite these advances, little is known about the functional mechanisms by which risk variants mediate disease susceptibility.

Increased understanding about the functional complexity of the genome has led to growing recognition about the likely role of non-sequence-based regulatory variation in health and disease. Building on the hypothesis that epigenomic dysregulation is important in the aetiology and progression of AD neuropathology^9^, we and others recently performed the first genome-scale cross-tissue analyses of DNA methylation in AD identifying robust DNA methylation differences associated with AD neuropathology across multiple independent human post-mortem brain cohorts^10, 11^. To date, however, no study has systematically examined other types of regulatory genomic modifications in AD. In this study, we focus on lysine H3K27 acetylation (H3K27ac), a robust mark of active enhancers and promoters that is strongly correlated with gene expression and transcription factor binding^12^. Interestingly, histone deacetylase (HDAC) inhibitors have been shown to ameliorate symptoms of cognitive decline and synaptic dysfunction in mouse models of AD^13, 14^ and are promising targets for novel human AD treatments^15, 16^. Despite this, investigations into global levels of histone acetylation in AD have thus far been inconclusive^17-20^ and no study has taken a genome-wide approach. In fact, few studies have systematically profiled H3K27ac across large numbers of samples in the context of complex disease, and optimal methods for these analyses are still being developed^21^.

In this study, we used chromatin immunoprecipitation combined with highly-parallel sequencing (ChIP-seq) to quantify levels of H3K27ac across the genome in post-mortem entorhinal cortex samples from AD patients and matched controls. We identify regulatory genomic signatures associated with AD, including variable H3K27ac across discrete regions annotated to genomic loci mechanistically implicated in the onset of both tau and amyloid pathology. This is the first study of variable H3K27ac yet undertaken for AD; in addition to identifying molecular pathways associated with AD neuropathology, we present a framework for genome-wide studies of this modification in complex disease.

## Results

### Genome-wide profiling of inter-individual variation in H3K27ac in the entorhinal cortex

We generated H3K27ac ChIP-seq data using post-mortem entorhinal cortex tissue dissected from 47 elderly individuals (average age = 77.43, SD = 9.66, range = 58-97) comprising both AD cases (n = 24, mean Braak stage = 6.00, SD = 0.00) and age-matched low pathology controls (n = 23, mean Braak stage = 1.30, SD = 1.11) **(Supplementary Table 1**). Raw H3K27ac ChIP-seq data is available for download from the Gene Expression Omnibus (GEO) (accession number GSE102538). After stringent quality control (QC) of the raw data (see **Methods**), we obtained an average of 30,032,623 (SD = 10,638,091) sequencing reads per sample, with no difference in read-depth between AD cases and controls (*P* = 0.93, **Supplementary Fig. 1**). This represents, to our knowledge, the most extensive analysis of H3K27ac in the human entorhinal cortex yet undertaken. Using combined data from all 47 samples (see **Methods**) we identified 182,065 high confidence H3K27ac peaks; these are distributed across all 24 chromosomes (**Supplementary Table 2**) spanning a mean length of 983bp (SD = 682bp) with a mean distance between neighbouring peaks of 15,536bp (SD = 116,040bp). We validated the identified peaks using two independent ChIP-seq datasets: first, we obtained locations for frontal cortex (BA9) and cerebellum H3K27ac peaks from a recent analysis of autism and control brains^21^; second, we downloaded peak profiles for multiple cell - and tissue-types from the NIH Epigenomics Roadmap Consortium^22^ (see **Methods**). As expected, there was a very high overlap between H3K27ac peaks called in other neocortical datasets and our ChIP-seq data, with a notably lower overlap with H3K27ac data from non-cortical tissues (**Supplementary Fig. 2**).

### AD-associated differential acetylation in the entorhinal cortex

We quantified read counts across every peak in each individual sample using *HTSeq*^23^ and employed a quasi-likelihood F test, implemented in the Bioconductor package *EdgeR*^24^, to test for differences in H3K27ac between AD cases and low pathology controls (see **Methods**). Our analysis model controlled for age at death and neuronal cell proportion estimates derived from DNA methylation data generated on the same samples using the *CETS* R package^25^ (**Supplementary Table 1**, **Supplementary Fig. 3**). Subsequent sensitivity analyses excluded an effect of sex on our results, but revealed widespread effects of cell-type proportion estimates and age at death highlighting the importance of including these covariates (**Supplementary Fig. 4**). A total of 4,162 (2.3%) of the 182,065 peaks were characterized by AD-associated differential acetylation at a false discovery rate (FDR) < 0.05 (**Fig. 1**), with a significant enrichment of hypo-acetylated AD-associated peaks (2,687 (1.5%)) compared to hyper-acetylated AD-associated peaks (1,475 (0.8%)) (*P* = 9.89E-80, exact binomial test) (**Fig. 1**). UCSC Genome Browser tracks showing H3K27ac levels in AD cases and controls, in addition to association statistics, across the genome can be accessed at https://tinyurl.com/y8f3tmoa. The ten top-ranked hyper - and hypo-acetylated peaks associated with AD are shown in **Table 1,**with a complete list given in **Supplementary Table 3** (hyperacetylated peaks) and **Supplementary Table 4** (hypoacetylated peaks). Peaks were subsequently annotated to genes using the Genomic Regions Enrichment of Annotations Tool (*GREAT*)^26^, which takes into account the strength of proximal and distal DNA-binding events. The most significant AD-associated hyperacetylated peak (chr13: 112101248-112102698; *P* = 2.04E-08; log fold change = 0.93) is annotated to both *SOX1* and *TEX29* on chromosome 13 (**Fig. 2**, **Table 1**). Of note, H3K27ac data from the Epigenomics Roadmap Consortium show that this region is characterized by brain-specific enhancer activity (**Supplementary Fig. 5**). The most significant AD-associated hypoacetylated peak (chr7: 64011549-64012825; *P* = 1.66E-08; log fold change = -0.86) is located within intron 1 of *ZNF680* on chromosome 7 (**Fig. 3**, **Supplementary Fig. 5**, **Table 1**). Global clustering of samples by normalized read counts across all hyper - and hypo - acetylated peaks (FDR < 0.05) indicated that, as expected, samples group primarily by disease status (**Fig. 4**). AD-associated differentially acetylated peaks (FDR < 0.05) are significantly longer (1295bp vs 975bp, *P* < 1.00E-50) and characterized by higher read - depths (2.56 log counts per million (CPM) vs 1.59 log CPM, *P* < 1.00E-50) than nonsignificant peaks (**Supplementary Fig. 6**). Of note, within AD-associated peaks, hypoacetylated peaks are significantly longer (1412bp vs 1081bp, *P* = 5.66E-31) and have higher read-depths (2.21 log CPM vs 1.77 log CPM, *P* = 2.69E-50) compared to hyperacetylated peaks. We used *RSAT*^27, 28^ to identify enriched transcription factor binding motifs located within AD-associated differentially acetylated peaks (see **Methods**), observing a significant enrichment of binding motifs for specificity protein 1 (Sp1) (*P* < 1.0E - 50) amongst AD-hyperacetylated peaks (FDR < 0.05). Of note, previous publications have reported dysregulated expression of Sp1 and its co-localization with neurofibrillary tangles in _AD_^29, 30^.

**Figure 1.**
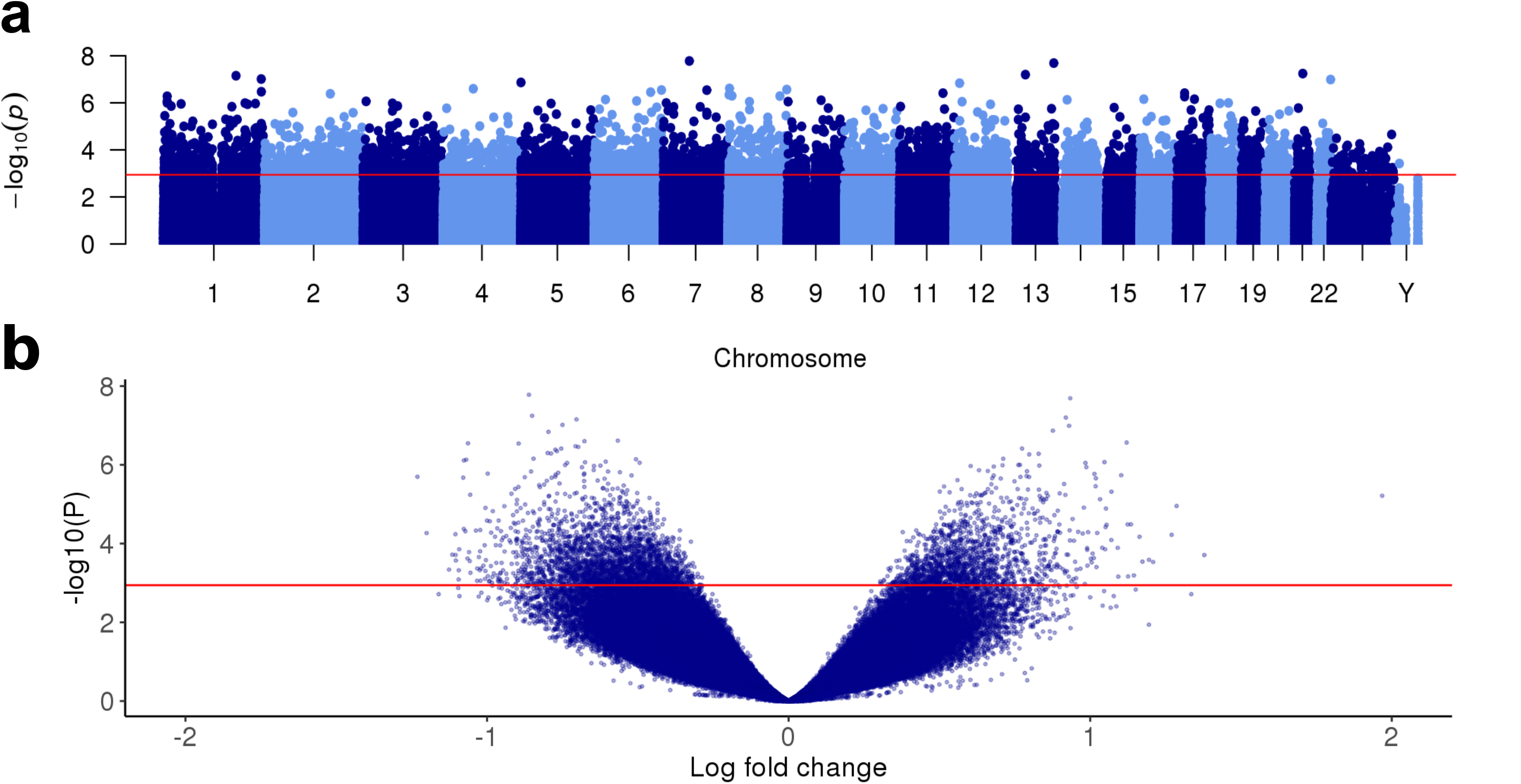
Variable H3K27ac associated with Alzheimer’s disease (AD) in the entorhinal cortex. (**a**) Manhattan plot showing the –log_10_ P value for differential H3K27ac against chromosomal location from the *EdgeR* quasi-likelihood F test, controlling for age and derived neuronal proportion. Variation in H3K27ac at 4,162 peaks was identified as being associated with AD (FDR < 0.05, red line). (**b**) Volcano plot showing the -log_10_ *P* value and log fold change for differential H3K27ac at each entorhinal cortex H3K27ac peak (FDR < 0.05, red line). Of the AD-associated peaks, 1,475 (35%) are hyperacetylated in AD and 2,687 (65%) are hypoacetylated in AD (lower H3K27ac) (*P* = 9.89E-80, exact binomial test).

**Table 1.** Differential H3K27ac associated with AD. Shown are the ten top-ranked hyper - and hypo-acetylated H3K27ac peaks, controlling for age at death and neuronal proportion estimates derived from DNA methylation data.

| Rank | CHR | Position (start – end) | P value | FDR | Log FC | Annotated genes |
| --- | --- | --- | --- | --- | --- | --- |
| Hyperacetylated peaks |  |  |  |  |  |  |
| 1 | 13 | 112101248-112102698 | 2.04E-08 | 0.002 | 0.93 | *SOX1, TEX29* |
| 2 | 13 | 42094789-42095919 | 6.31E-08 | 0.003 | 0.92 | *RGCC, VWA8* |
| 3 | 22 | 50342521-50343567 | 1.02E-07 | 0.003 | 0.93 | *PIM3, CRELD2* |
| 4 | 5 | 640598-642071 | 1.36E-07 | 0.003 | 0.88 | *CEP72, TPPP* |
| 5 | 8 | 145180336-145181125 | 2.72E-07 | 0.004 | 1.12 | *FAM203A, MAF1* |
| 6 | 17 | 19665361-19666514 | 3.86E-07 | 0.004 | 0.77 | *ALDH3A1, ULK2* |
| 7 | 1 | 9392591-9393233 | 5.25E-07 | 0.004 | 0.83 | *SLC25A33, SPSB1* |
| 8 | 17 | 19619421-19620832 | 5.43E-07 | 0.004 | 0.80 | *SLC47A2* |
| 9 | 17 | 43925717-43927482 | 7.01E-07 | 0.005 | 0.71 | *MAPT, SPPL2C* |
| 10 | 1 | 9341867-9342320 | 8.55E-07 | 0.005 | 1.05 | *SPSB1, H6PD* |
| Hypoacetylated peaks |  |  |  |  |  |  |
| 1 | 7 | 64011549-64012825 | 1.66E-08 | 0.002 | -0.86 | *ZNF680, ZNF736* |
| 2 | 21 | 29827289-29828201 | 5.70E-08 | 0.003 | -0.85 | *N6AMT1* |
| 3 | 1 | 179175226-179176637 | 7.03E-08 | 0.003 | -0.70 | *ABL2, TOR3A* |
| 4 | 1 | 241397411-241399621 | 9.73E-08 | 0.003 | -0.75 | *GREM2, RGS7* |
| 5 | 12 | 13627258-13629064 | 1.46E-07 | 0.003 | -0.80 | *EMP1, GRIN2B* |
| 6 | 8 | 3964265-3966191 | 2.44E-07 | 0.004 | -0.57 | *CSMD1* |
| 7 | 4 | 74088063-74089559 | 2.52E-07 | 0.004 | -0.68 | *COX18, ANKRD17* |
| 8 | 6 | 166401119-166402753 | 2.85E-07 | 0.004 | -1.06 | *SDIM1, T* |
| 9 | 7 | 107111795-107113029 | 2.88E-07 | 0.004 | -0.90 | *DUS4L, GPR22* |
| 10 | 1 | 241694436-241695782 | 3.37E-07 | 0.004 | -0.71 | *KMO* |

**Figure 2.**
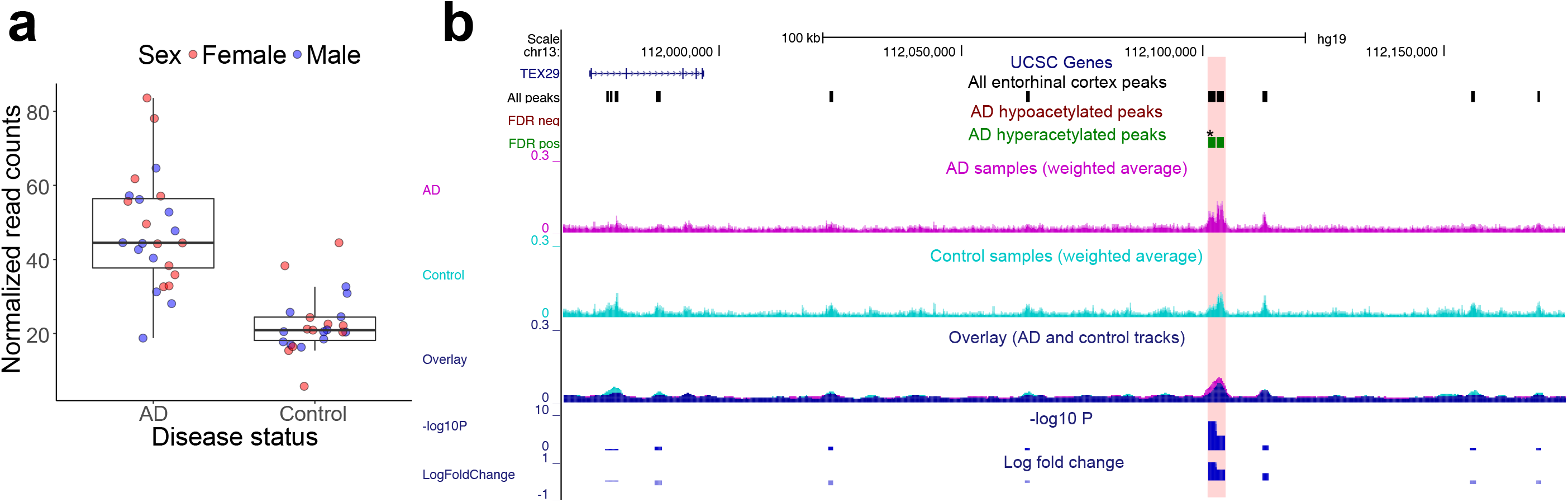
The top-ranked AD-associated hyperacetylated peak is annotated to *SOX1* and *TEX29* on chromosome 13. Shown are (**a**) normalized read counts and (**b**) a regional track of H3K27ac ChIP-seq data showing weighted average H3K27ac profiles of cases (magenta), controls (turquoise) as well as an overlay track, highlighting the acetylation differences. (**a**) The most significant AD-hyperacetylated peak is characterized by a consistent increase in H3K27ac in patients (*P* = 2.04E-08, log fold change = 0.93). (**b**) This peak is located on chromosome 13 and annotated to both *SOX1* and *TEX29*. Also shown is the location of all entorhinal cortex H3K27ac peaks in this region, hyper - and hypoacetylated peaks (FDR < 0.05), as well as the -log 10 *P* value and log fold change of normalized read count differences for each peak.

**Figure 3.**
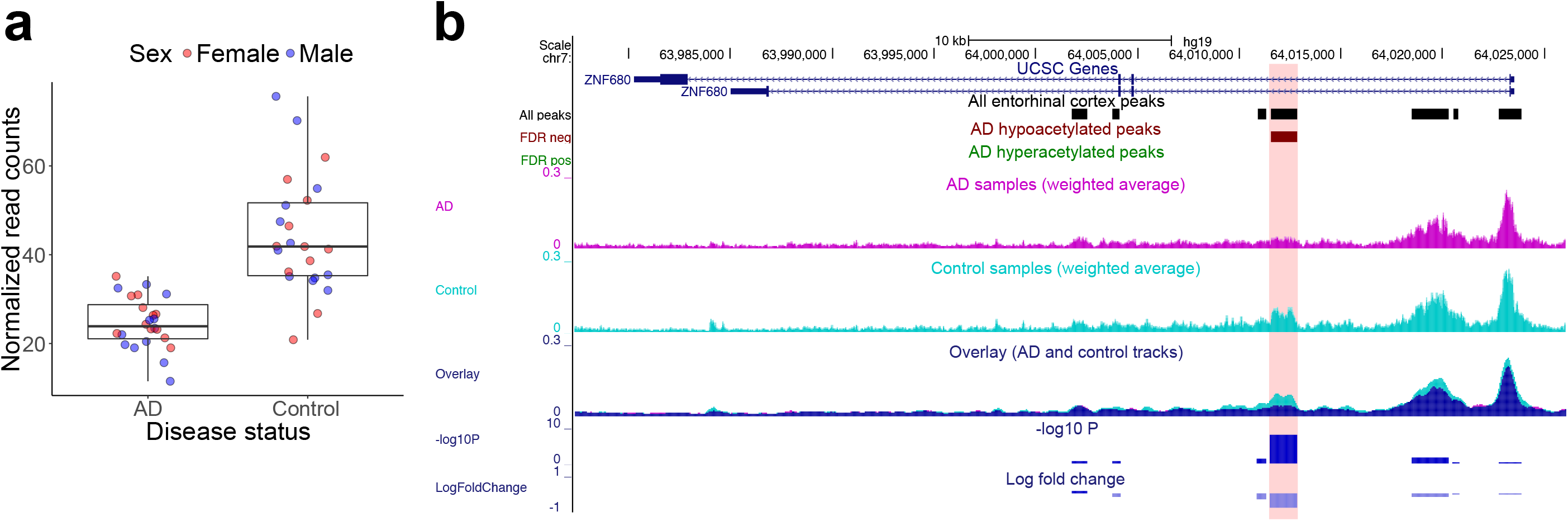
The top-ranked AD-associated hypoacetylated peak is located in intron 1 of *ZNF680* on chromosome 7. Shown are (**a**) normalized read counts and (**b**) a regional track of H3K27ac ChIP-seq data showing weighted average H3K27ac profiles of cases (magenta), controls (turquoise) as well as an overlay track, highlighting the acetylation differences. (**a**) The most significant AD-hypoacetylated peak (*P* = 1.66E-08) is characterized by a consistent decrease in H3K27ac in cases (log fold change = -0.86). (**b**) This peak is located in intron 1 of *ZNF680* on chromosome 7. Also shown is the location of all entorhinal cortex H3K27ac peaks in this region, hyper - and hypoacetylated peaks (FDR < 0.05), as well as the -log 10 *P* value and log fold change of normalized read count differences for each peak.

**Figure 4.**
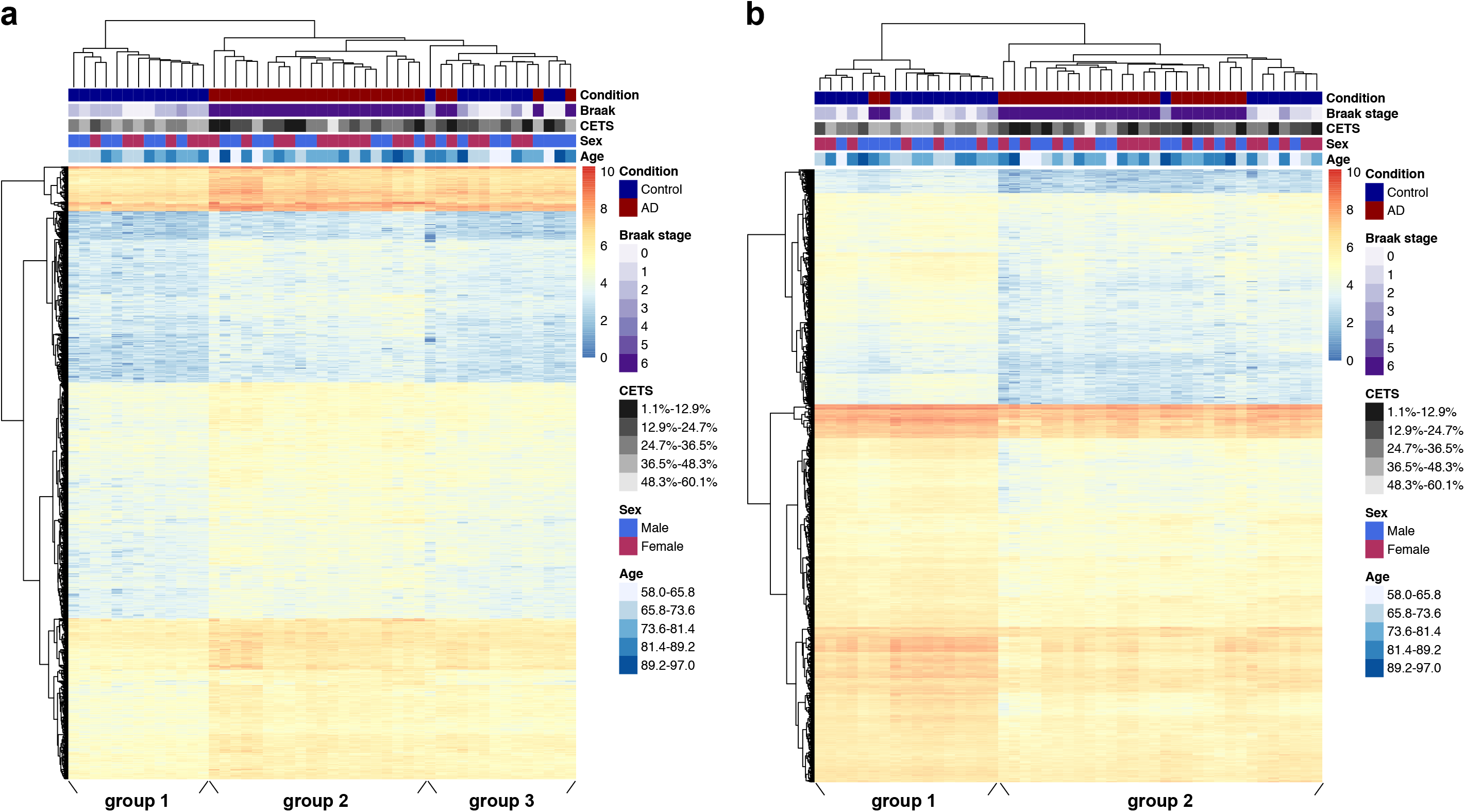
Clustering of AD and low pathology control samples by H3K27ac levels at differentially acetylated peaks. (**a**) A heatmap, clustering samples by normalized read counts in all 1,475 significant AD hyperacetylated peaks (FDR < 0.05), generates three distinct groups: one comprised of controls only (group 1), a pure group of cases (group 2), and a mixed group containing both cases and controls (group 3). Controls grouped together with cases in group 3 are characterized by significantly decreased neuronal proportion estimates, compared to those in the pure control group 1 (*P* = 7.10E-04, t-test). (**b**) A heatmap, clustering samples by all 2,687 significant AD hypoacetylated peaks (FDR < 0.05), divides the samples into two main groups: group 1 is composed mainly of controls, whereas group 2 contains more cases than controls. Interestingly, controls classified into group 2 are characterized by lower neuronal proportion estimates than those in group 1 (*P* = 0.004, t-test). The clustering defined by hyper - or hypoacetylated peaks is not significantly associated with sex (*P* > 0.05, chi-square test) or age at death (*P* > 0.05, t-test).

### Increased H3K27ac is observed in regulatory regions annotated to genes previously implicated in both tau and amyloid neuropathology

One of the top-ranked AD-associated hyper-acetylated peaks is located proximal to the gene encoding microtubule associated protein tau (MAPT) (chr17: 43925717-43927482; *P* = 7.01E-07; log fold change = 0.71; **Table 1**), which is widely expressed in the nervous system where it functions to promote microtubule assembly and stability. Tau is believed to play a key role in AD neuropathology, with hyperphosphorylation of the tau protein precipitating the neurofibrillary tangles associated with the pathogenesis of AD^31, 32^. Closer inspection of the region around this AD-associated peak highlighted an extended cluster of six hyper - acetylated H3K27ac peaks (FDR < 0.05) spanning 36kb (chr17: 43925717 - 43961546) located within a *MAPT* antisense transcript (*MAPT_AS1*) ~10kb upstream of the *MAPT* transcription start site (**Fig. 5**; **Supplementary Table 5**). H3K27ac ChIP-seq data from the NIH Epigenomics Roadmap Consortium show that this region is characterized by CNS - related H3K27ac signatures (**Fig. 5**), with *ChromHMM*^33^ identifying the region as an active chromatin domain in brain comprised of enhancers and blocks of weak transcription (**Supplementary Fig. 7**). Strikingly, AD-associated differentially-acetylated peaks were also found in the vicinity of other genes known to play a direct mechanistic role in AD. We identified a significantly hypoacetylated peak (chr21: 27160993 - 27161475; *P* = 3.94E-04; log fold change = -0.72) on chromosome 21, located ~100kb downstream of the amyloid precursor protein gene (*APP*), which encodes the precursor molecule to Ab, the main component of amyloid plaques^34-36^ (**Supplementary Fig. 8**). We also identified significant hyperacetylation in the vicinity of the presenilin genes *PSEN1* and *PSEN2*, which encode integral components of the gamma secretase complex and play a key role in generation of Aβ from APP^37^. In *PSEN1* we found significantly elevated H3K27ac across a peak within intron 6 (chr14: 73656445 - 73656860; *P* = 3.44E-04; log fold change = 0.68; **Supplementary Fig. 9**). In *PSEN2* we identified consistent hyperacetylation in AD cases across nine H3K27ac peaks (FDR < 0.05) spanning a ~57 kb region upstream of the transcription start-site (chr1: 226957424 - 227014019; **Fig. 6, Supplementary Fig. 7** and **Supplementary Table 6**). Of note, highly-penetrant mutations in *APP*, *PSEN1*, and *PSEN2* are associated with familial forms of early-onset AD^38^. The identification of altered regulation of these loci in late-onset sporadic AD brain further supports a key role for altered amyloid processing in the onset of neuropathology.

**Figure 5.**
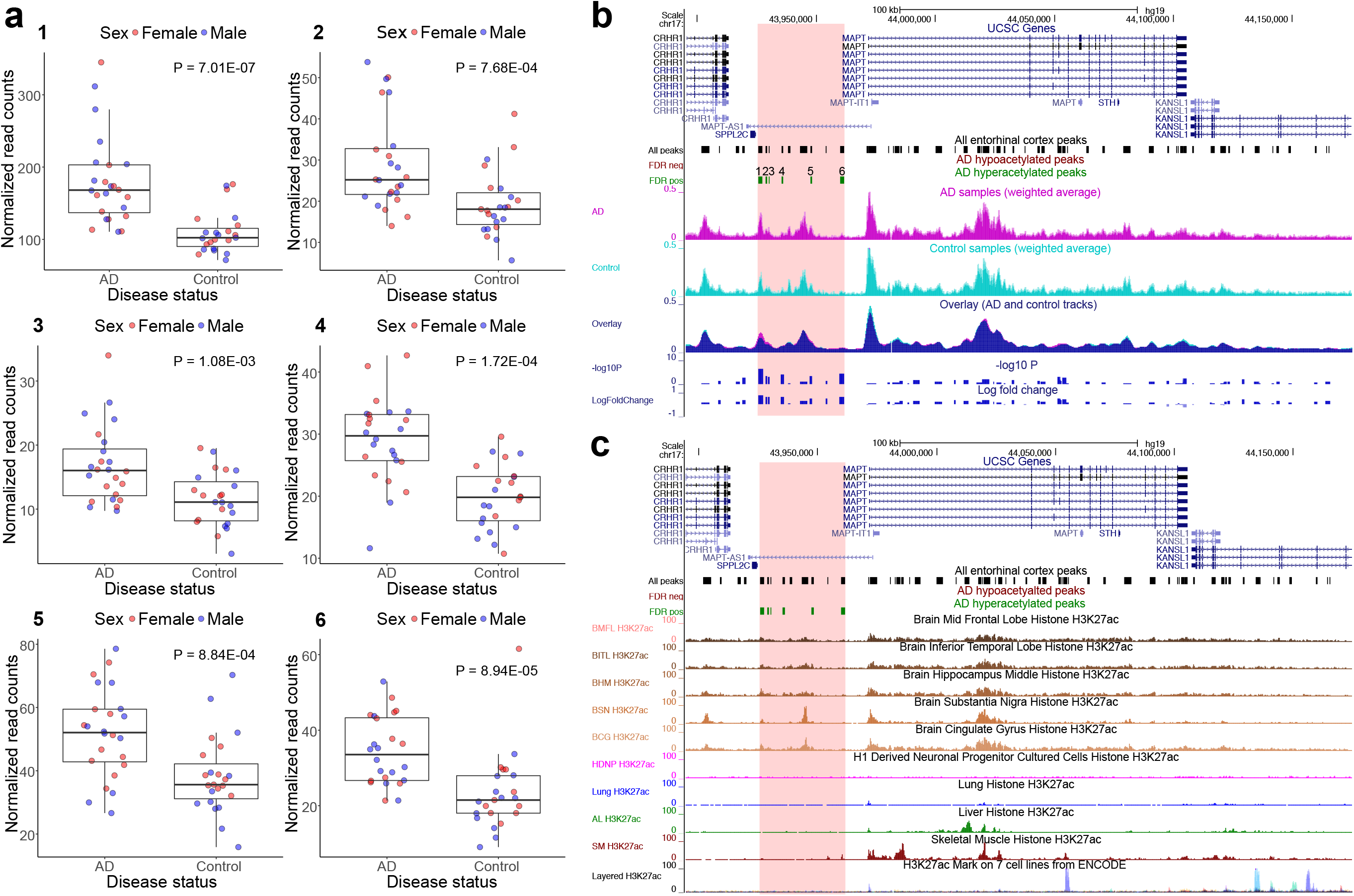
A region annotated to *MAPT* spanning six H3K27ac peaks is characterized by significant hyperacetylation in AD. A cluster of nine H3K27ac peaks was identified on chromosome 17. All nine peaks are hyperacetylated in cases (mean log fold change = 0.46; Supplementary Table 5). (**a**) For six of the nine peaks this increase in H3K27ac associated with AD is significant (FDR < 0.05). (**b**) The whole region is located 10kb upstream of *MAPT* (**c**) and is characterized by brain specific acetylation profiles. The boundaries of the significantly differentially acetylated peak region are highlighted in red.

**Figure 6.**
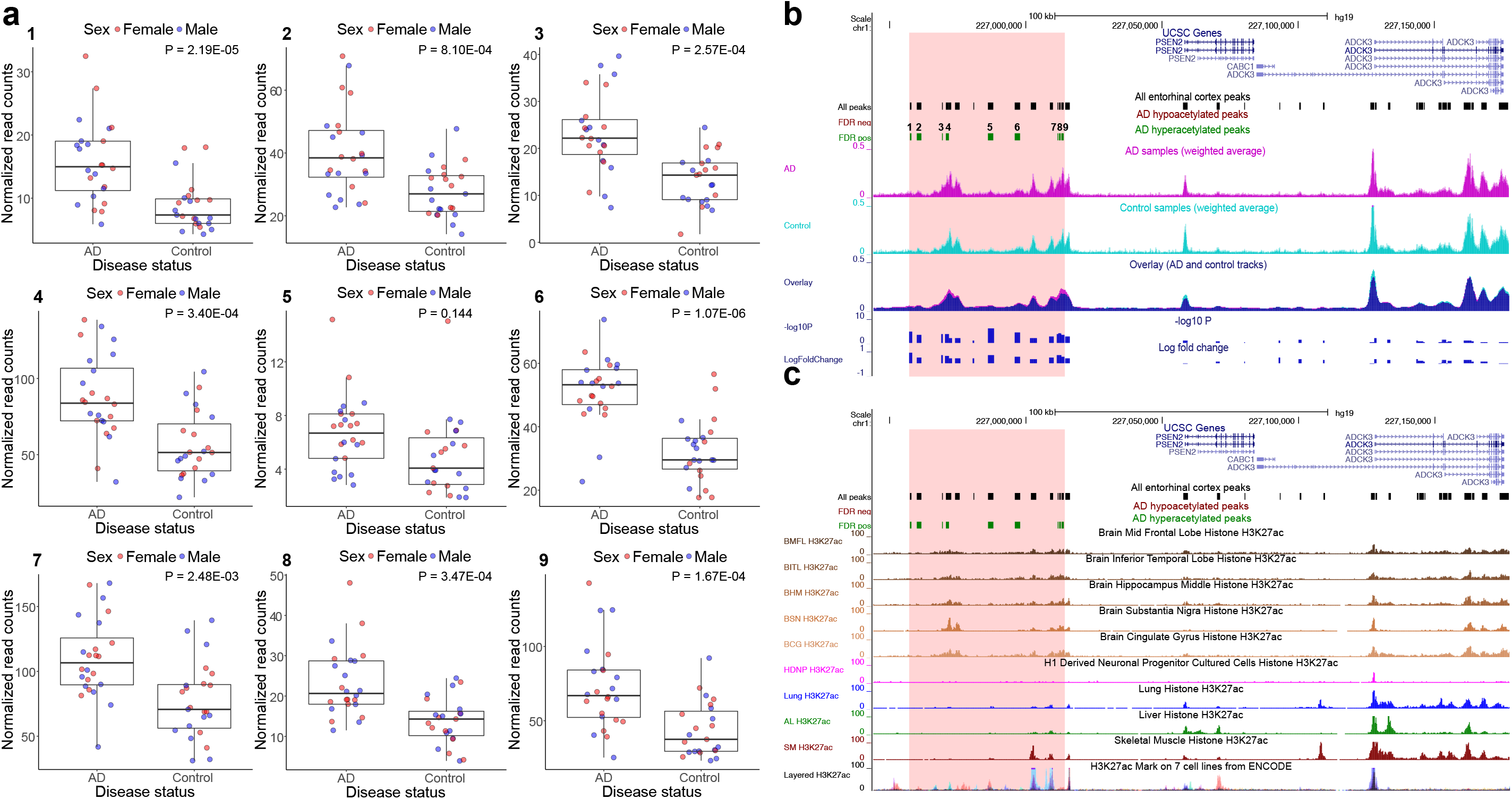
A region annotated to *PSEN2* spanning nine H3K27ac peaks is characterized by significant hyperacetylation in AD. A cluster of 14 H3K27ac peaks was identified on chromosome 1. All 14 peaks are hyperacetylated in cases (mean log fold change = 0.52; Supplementary Table 6). (**a**) For nine of the 14 peaks this increase in H3K27ac associated with AD is significant (FDR < 0.05). (**b**) The whole region is located 44kb upstream of *PSEN2* (**c**) and is characterized by predominantly brain-specific acetylation profiles. The boundaries of the significantly differentially acetylated peak region are highlighted in red.

### Specific differentially-acetylated peaks also overlap known AD GWAS regions

Using data from a large GWAS meta-analysis of AD^8^ we tested for instances where there is an overlap between AD-associated differential H3K27ac and genomic regions harbouring risk variants. Briefly, we defined linkage-disequilibrium (LD) blocks around the genome-wide significant (*P* < 5.0E-08) GWAS variants identified by the stage 1 meta-analysis by Lambert and colleagues^8^ (**Supplementary Table 7**), which contained a total of 292 overlapping entorhinal cortex H2K27ac peaks (see **Methods**). Two of the 11 GWAS LD blocks contained significant AD-associated H3K27ac peaks (FDR < 0.05), although there was no overall enrichment of AD-associated differential acetylation at the 292 peaks (*P* = 0.364, Wilcoxon rank-sum test). Two peaks of AD-associated hyperacetylation were located within a GWAS region on chromosome 1, mapping to the gene body of *CR1* (chr1: 207753457-207753813; *P* = 1.15E-06; log fold change = 0.99 and chr1: 207754916-207756572; *P* = 5.40E-04; log fold change = 0.56; **Supplementary Fig. 10**). *CR1* encodes a transmembrane glycoprotein expressed in microglia with a role in the innate immune system, promoting phagocytosis of immune complexes and cellular debris, in addition to Aβ^39-41^. Two other AD-associated differentially acetylated peaks were found to be located within a GWAS region on chromosome 19, including a hyperacetylated peak (chr19:45394441-45395396; *P* = 2.13E-04; log fold change = 0.48) mapping to the gene body of *TOMM40* in the immediate vicinity of *APOE* (**Supplementary Fig. 11**). Another H3K27ac peak in this LD block was significantly hypoacetylated in AD (chr19: 45639588-45641733; *P* = 7.65E-04; log fold change = -0.33), mapping to intron 1 of *PPP1R37*.

### AD-associated differentially-acetylated peaks are enriched for functional processes related to neuropathology

We used *GREAT*^26^ to calculate statistical enrichments for ontological annotations amongst our AD-associated peaks, interrogating gene ontologies for molecular function and biological processes^42^, human diseases^43^ as well as human phenotypes^44^ (**Supplementary Fig. 12** and **Supplementary Tables 8-15**). Multiple ontological categories associated with AD progression and pathology were identified as being enriched (FDR < 0.05) amongst hyperacetylated peaks, including “lipoprotein particle binding”^7, 45^ (*P* = 1.10E-06), “beta - amyloid metabolic process”^31^ (*P* = 4.94E-08), “neurofibrillary tangles”^32^ (*P* = 3.93E-05), “response to hypoxia”^46, 47^ (*P* = 3.17E-14), “neuronal loss in central nervous system”^2^ (*P* = 4.93E-04), and “Pick’s disease” (*P* = 2.93E-07), a form of fronto-temporal dementia also characterized by tau pathology^32, 48^. Amongst hypoacetylated peaks we observed an enrichment of categories related to neurotransmitter-functions, including “GABA receptor activity”^49^ (*P* = 2.70E-07) as well as categories related to neuronal transmission and synapses, such as “protein location to synapse” (*P* = 7.86E-09).

### Integrative analysis of DNA and histone modifications reveal unique distributions of DNA modifications across regions of differential acetylation

Our previous work identified cortex-specific variation in DNA methylation (5mC) robustly associated with AD pathology^10, 11^. We were therefore interested in exploring the relationship between H3K27ac and both 5mC and another DNA modification – DNA hydroxymethylation (5hmC), which is enriched in the brain and believed to play an important role in neuronal function, learning and memory^50-52^ - in AD. Both modifications were profiled using DNA isolated from the same entorhinal cortex samples using oxidative bisulfite (oxBS) conversion in conjunction with the Illumina 450K HumanMethylation array (“450K array”) (see **Methods**). 6,838 loci mapped to within 1kb of differentially acetylated peaks (FDR < 0.5; mapping to 1,649 unique peaks). We tested differential 5mC and 5hmC associated with AD at these probes, controlling for age at death and cell-type proportion estimates. None of the differences in 5mC (minimum *P* = 2.47E-03) or 5hmC (minimum *P* = 1.53E-03) were significant when correcting for multiple testing (n = 6,838 tests; *P* < 7.31E-05), indicating that there is little direct overlap in AD-associated variation in H3K27ac and DNA modifications. Furthermore, comparing effect sizes at these 6,838 peak–probe pairs identified no evidence for a correlation between AD-associated H3K27ac and 5mC differences (r = 0.009, *P* = 0.443; **Supplementary Fig. 13**) with a small, but significant, negative correlation for 5hmC (r = -0.045, *P* = 1.63E-04; **Supplementary Fig. 13**). As expected, both DNA modifications are significantly lower in the vicinity of H3K27ac peaks compared to the genome-wide 450K array background (5mC: *P* < 1.00E-50, beta difference = 12%, **Supplementary Fig. 13**; 5hmC: *P* = 3.61E-30, beta difference = 0.16%; **Supplementary Fig. 13**), consistent with H3K27ac being localized at active enhancers and promoters.

## Discussion

We quantified H3K27ac across the genome in post-mortem entorhinal cortex tissue samples, identifying widespread AD-associated acetylomic variation. Strikingly, differentially acetylated peaks were identified in the vicinity of genes implicated in both tau and amyloid neuropathology as well as genomic regions containing variants associated with sporadic late-onset AD. This is the first study of variable H3K27ac yet undertaken for AD; in addition to identifying molecular pathways associated with AD neuropathology, we introduce a framework for genome-wide studies of this modification in complex disease.

Given its close relationship with transcriptional activation, for example via the mediation of transcription factor binding, the identification of AD-associated variation in H3K27ac highlights potential novel regulatory genomic pathways involved in disease etiology. We find widespread alterations in H3K27ac associated with AD, including in the vicinity of several genes known to be directly involved in the progression of Abβand tau pathology^31, 53^ (*APP*, *PSEN1*, *PSEN2*, *MAPT*), supporting the notion that dysregulation of both pathways is involved in the onset of AD. Interestingly, although our study assessed brains from donors affected by sporadic late-onset AD, we identify widespread altered H3K27ac in the vicinity of genes implicated in familial early-onset AD. This indicates that these two forms of the disease may share common pathogenic pathways and mechanisms. Given that histone - acetylation modifiers are amongst the most promising target pharmacological treatments of AD^15, 54^, the identification of altered H3K27ac in AD is important, giving clues as to which genes and pathways may be involved.

Our study has a number of limitations, which should be considered when interpreting these results. First, we undertook ChIP-seq using bulk entorhinal cortex samples comprising a mix of neuronal and non-neuronal cell-types. This is an important limitation in epigenomic studies of a disease characterized by cortical neuronal loss. However, we were able to control for variation in neuronal proportions in our samples by deriving neuronal proportion estimates for each sample using DNA methylation data generated on the same tissue samples^25^. Second, our cross-sectional analysis of post-mortem brain tissue makes direct causal inference difficult, and it is likely that many of the changes in H3K27ac we observe result from the AD pathology itself. In this regard, however, it is interesting that we see disease-associated H3K27ac in the vicinity of genes causally implicated in familial forms of AD. Third, we have assessed a relatively small number of samples. In this light, it is notable that we identify substantial differences between AD cases and controls, with disease - associated regulatory variation in genes and functional pathways known to play a role in the onset and progression of neuropathology. The clear clustering between patients and controls at our differentially acetylated peaks suggests that, despite a complex and heterogeneous etiology, AD may be characterized by a common molecular pathology in the entorhinal cortex, reflecting neuropathological analyses. Fourth, chromatin architecture and transcriptional regulation is influenced by a multitude of epigenetic mechanisms. Although profiling H3K27ac can provide relatively robust information about transcriptional activity, it represents only one of perhaps ~100 post-translational modifications occurring at > 60 histone amino-acid residues regulating genomic function.

In summary, we provide compelling evidence for widespread acetylomic dysregulation in the entorhinal cortex in AD. Our data suggest that regulatory variation at multiple loci, including in the vicinity of several known AD risk genes – *APP*, *CR1*, *MAPT*, *PSEN1*, *PSEN2* and *TOMM40* – is robustly associated with disease, supporting the notion of common molecular pathways in both familial and sporadic AD. In addition to identifying molecular pathways associated with AD neuropathology, we present a framework for genome-wide studies of histone modifications in complex disease, integrating our data with results obtained from genome-wide association studies as well as other epigenetic marks profiled on the same samples.

## Online Methods

### Samples

Post-mortem brain samples from 54 individuals - 27 with advanced AD neuropathology and 27 neuropathology-free brain samples - were provided by the MRC London Neurodegenerative Disease Brain Bank (http://www.kcl.ac.uk/ioppn/depts/cn/research/MRC-London-Neurodegenerative-Diseases-Brain-Bank/MRC-London-Neurodegenerative-Diseases-Brain-Bank.aspx). Ethical approval for the study was provided by the NHS South East London REC 3. Samples for this ChIP-seq study were selected from a larger collection of post-mortem entorhinal cortex (Brodmann area (BA) 28/34) samples, based on Braak staging, a standardized measure of neurofibrillary tangle burden determined at autopsy^55^. We prioritized cases with high Braak staging and controls with lower Braak scores (**Supplementary Table 1**). Subjects were approached in life for written consent for brain banking, and all tissue donations were collected and stored following legal and ethical guidelines (NHS reference number 08/MRE09/38; the HTA license number for the LBBND brain bank is 12293). All samples were dissected by trained specialists, snap-frozen and stored at -80 °C. A detailed list of demographic and sample data for each individual included in the final analyses is provided in **Supplementary Table 1**.

### Chromatin immunoprecipitation

Chromatin immunoprecipitation was performed using the iDeal ChIP-Seq kit for Histones (Cat# C01010051, Diagenode, Seraing, Belgium) as detailed below, using the standard kit components unless otherwise stated. 30 mg of entorhinal cortex tissue was homogenized with a dounce homogenizer in 1 mL ice-cold phosphate buffered saline (PBS) buffer with protease inhibitor cocktail (PIC). The suspension was centrifuged at 4,000 rpm for 5 minutes at 4°C, discarding the supernatant. The pellets were resuspended in 1 mL PBS containing 1% formaldehyde, rotating at room temperature for 8 minutes. The cross-linking process was terminated by adding 100 μL glycine solution, followed by 5 minutes of rotation. After 5 minutes of centrifugation at 4,000 rpm and 4°C, the pellet was washed twice with ice-cold PBS (suspending the pellet in 1 mL PBS with PIC, centrifuging for 5 minutes at 4,000 rpm and 4°C, and discarding the supernatant), then lysed in 10 mL ice-cold lysis buffer iL1 and iL2, sequentially (re-suspending the pellet in 10 mL lysis buffer, mixing gently for 10 minutes at 4°C, centrifuging for 5 minutes at 4,000 rpm and 4°C, and discarding the supernatant). The cross-linked lysate was suspended in 1.8 mL shearing buffer iS1 containing PIC and sonicated in aliquots of 300 μL for 10 cycles (30 seconds on/off each cycle) on a Bioruptor Pico (Diagenode, Seraing, Belgium). After shearing, samples were transferred to 1.5 mL microcentrifuge tubes and centrifuged at 14,000 rpm for 10 minutes, collecting the supernatant, containing the soluble sheared chromatin with fragments of an average size range of 100-1000bp as visualized by agarose gel electrophoresis (**Supplementary Fig. 14**).

Immunoprecipitation was performed on the SX-8G IP-Star robot (Diagenode, Seraing, Belgium), following the manufacturer’s protocol. All samples were immunoprecipitated with H3K27ac polyclonal antibody (Diagenode, Cat #C15410196, Seraing, Belgium). In addition, a randomly selected subgroup of 12 samples – 6 cases and 6 controls – were immunoprecipitated with rabbit IgG antibody (iDeal ChIP-seq kit) as negative control. 1-1.5 μL of H3K27ac or IgG antibody were first mixed with 98.5-99 μL ChIP buffer iC1, 0.5 μL PIC and 4 μL of 5% bovine serum albumin (BSA), which was incubated with magnetic beads for 3 hours at 4°C. Next the antibody conjugate was added to 180 μL chromatin for overnight (15h) immunoprecipitation at 4°C in an immunoprecipitation mix also containing 20 μL ChIP buffer iC1, 1 μL PIC and 4 μL of 5% BSA. After immunoprecipitation, the beads were resuspended in in 100 μL elution buffer iE1, to which 4 μL elution buffer iE2 was added. Cross-link reversal was performed on a PCR thermoblock for 4 hours at 65°C. DNA was extracted using Micro ChIP DiaPure columns (Diagenode, Cat No C03040001, Seraing, Belgium) according to the manufacturers protocol, eluting the DNA from the column matrix in 30 μL DNA elution buffer (MicroChIP DiaPure columns; Diagenode, Cat #C03040001, Denville, NJ, USA). Quantitative PCR, using 1% input DNA, was used to confirm specific enrichment of H3K27ac at positive control genes (*IEF4A2* and *GAPDH*) but not at negative control genes (*MBex2* and *TSH2b*; all primers were provided by Diagenode).

### Sequencing

Libraries were prepared using the MicroPlex Library Preparation kit v2 (Diagenode, Cat #C05010013, Seraing, Belgium) on a SX-8G IP-Star robot according to the manufacturer’s protocol. DNA concentrations were measured with Qubit dsDNA HS Assay Kits (Invitrogen, Cat# Q32851, Carlsbad, CA, USA) on the Qubit 2.0 Fluorometer (Invitrogen) and library fragment profiles generated on the Agilent 2100 BioAnalyzer using Agilent High Sensitivity DNA kits (Agilent Technologies, Cat# 5067-4626, Santa Clara, CA, USA). Following our stringent quality-control filtering, 7 samples were excluded from sequencing based on poor qPCR results after immunoprecipitation or low library concentration. The remaining 47 samples (from 24 cases and 23 controls) were sequenced on an Illumina HiSeq-2500 using single-end sequencing and a read length of 50bp. ChIP-seq data are available to download from GEO (accession number GSE102538).

### Data pre-processing and quality control

Global sample anomalies were ruled out using *fastqc*^56^ summary measures. All fastq files were aligned to the *Homo sapiens* reference genome (hg19, Broad Institute) using *Bowtie*^57^. The output SAM files were converted to binary (BAM) format. All BAM files were sorted and indexed using *samtools*^58^. PCR duplicates were removed using *Picard* (http://broadinstitute.github.io/picard/). *Samtools* was used to additionally remove nonuniquely mapped reads as well as reads with a sequencing quality score q < 30. Final read counts after QC for all 47 samples are shown in **Supplementary Fig. 1.** On average, we obtained 30,032,623 reads per sample (SD= 10,638,091; range=10,910,000-53,770,000) and individual read counts did not associate with disease status (*P* = 0.93).

### Peak calling and read counts

All filtered BAM files were merged into one grouped file and converted to *tagAlign* format using *bedtools*^59^. Peaks were called on this merged file using *MACS2*^60^, keeping all duplicates, since duplicates were removed from each sample previously and any remaining duplicates would result from the same read occurring in more than one sample. From the resulting peaks those located in unmapped contigs and mitochondrial DNA were filtered out as well as peaks that did not meet a significance threshold of *P* < 1.0E-07 for peak calling. The bed file of peaks was converted to gff format using *awk* and *R*, and reads for each individual sample were generated using *HTSeq*^23^. Final filtering was performed using the Bioconductor package *EdgeR*^*24*^, excluding peaks with fewer than 2 samples showing at least 1 read per million, resulting in a total of 182,065 peaks to be tested. Principal components analysis (PCA) in *R* using *DESeq2*^61^ confirmed that the epigenetically predicted gender was identical to the recorded one (**Supplementary Fig. 15**), with load on the first two principal components not related to disease status.

### Peak validation

We validated the 182,065 union peaks in two ways. First, we obtained the locations of H3K27ac peaks called in the cortex (BA9) and cerebellum from a recent paper by Sun and colleagues^21^. Second, we downloaded H3K27ac profiles produced by the NIH Roadmap Epigenomics Consortium^22^ from the Gene Expression Omnibus (GEO; https://www.ncbi.nlm.nih.gov/geo) for multiple cell-/tissue-types including several brain regions (mid frontal lobe (GSM773015), inferior temporal gyrus (GSM772995), middle hippocampus (GSM773020), substantia nigra (GSM997258), cingulate gyrus (GSM773011), H1-derived neuronal progenitor cells (HDNPs, GSM753429), lung (GSM906395), liver (GSM1112808) and skeletal muscle (GSM916064)). The downloaded files were in bed format, on which we performed peak calling using *MACS2* and the same specifications as described for our own samples, discounting any duplicate reads. We calculated the overlap between each peak set and our peaks by quantifying the percentage of peaks from the external sample overlapping our peaks using the Bioconductor package *GenomicRanges*^62^.

### Differential peak calling

We used the quasi-likelihood F test^63^ in *EdgeR*^24^ to analyse peak differences between AD - cases and controls, allowing us to correct for potential confounders in the analysis of differential peaks. Our analyses accounted for additional phenotypic variation across the samples, including age at death and neuronal proportion estimates based on DNA methylation profiles from the Illumina 450K HumanMethylation Array from the same samples, which were calculated using the *CETS* R package^25^. To test for the effects of these two covariates and sex we ran three further analyses, omitting age at death and CETS and adding sex as a covariate to our main model. We imputed the median CETS estimate for one individual with missing DNA methylation data. To include age at death and CETS estimates in the *EdgeR* differential peak calling method, these variables were converted to five-level factors using the R function *cut()*. The distribution of the age and CETS variable (including the imputed individual) with the respective bins of the factor variables are shown in **Supplementary Fig. 3**. We next calculated normalization factors based on sample specific library compositions and estimated both sample and peak-specific dispersions, specifically for a generalized linear model controlling for factorized CETS estimates and age at death. The quasi-likelihood F-test was conducted after fitting a quasi-likelihood model^63^ using the *glmQLFit()* and *glmQLFTest()* functions respectively. Effect sizes are reported as log fold change, a standard measure for quantifying sequencing read count differences between different conditions. Log fold change refers to the log2-transformed ratio of normalized read counts between cases and controls, with positive values indicating higher normalized read counts in AD samples. The *bedtools* program *genomecov* was used to generate coverage value scaled by library size and the number of samples per group, for each sample. These were then joined using *unionbedg* and summed using a *Perl* script to produce a weighted mean for each variable sized interval defined by read overlaps.

### Genomic annotation and enrichment analyses

Peaks were annotated to genes using the *Genomic Region Enrichment and Annotation Tools* (*GREAT*)^26^. In addition, we performed enrichment analyses calculate statistical enrichments for ontological annotation (gene ontologies for molecular function and biological processes^42^, human diseases^43^ as well as human phenotypes^44^). Functional enrichment analyses were conducted for significantly hyper - and hypoacetylated peaks (FDR < 0.05) separately, using the basal plus extension option. Significance in the enrichment test is based on a hypergeometric test of genes annotated to the test set (hyper-/ hypoacetylated peaks) compared to the background set of genes annotated to all 182,065 peaks called across all samples. Results presented in **Supplementary Fig. 11** are restricted to the top five non-redundant enrichments (separated by at least two nodes in the local directed acyclic graph visualizing the hierarchy of enriched terms from a single ontology) associated with at least three genes in the test set for the ontology categories biological process, molecular function, and disease ontology and we show full enrichments across all categories in **Supplementary Tables 8-15**.

### Motif enrichment analysis

Motif analysis was performed using the *Regulatory Sequence Analysis Tools suite* (*RSAT*)^27, 28^, available at http://rsat.sb-roscoff.fr. Peak sequences were reduced to 500bp on each side of the peak centre, and motif discovery was conducted on 6 and 7mer oligonucleotides, comparing the statistically enriched sequences with known transcription factor motifs from *JASPAR*^64^ (core nonredundant vertebrates) and *Homer*^65^ (Human TF motifs). Enrichments were computed relative to the background peak sequences (n = 182,065 peaks) for significantly hyper - and hypoacetylated peaks (FDR < 0.05).

### Analysis of differential H3K27ac across AD regions from genome-wide association studies (GWAS)

The summary statistics for the stage 1 GWAS from Lambert and colleagues^8^ were downloaded from http://web.pasteur-lille.fr/en/recherche/u744/igap/igap_download.php. These results were clumped (p1 = 1e-4; p2 = 1e-4, r2 =0.1, window = 3000kb) using *plink*^66^, which collapses multiple correlated signals (due to linkage disequilibrium (LD)) into regions which represent independent signals. LD relationships were inferred from a reference GWAS dataset (Phase 1) from another study^67^. Neighbouring regions located within 250kb of each other on the same chromosome were subsequently merged. After clumping, each region was assigned the minimum *P* value for all SNPs contained in the region (from Lambert et al), and regions were then filtered to the genome-wide significance threshold (*P* < 5.0E-08). This yielded 11 LD blocks for the genome-wide significant findings from Lambert et al., which were then overlapped with our AD-associated differentially acetylated peaks using the Bioconductor package *GenomicRanges*^62^.

### Integrative analysis with DNA methylation and hydroxymethylation

DNA methylation and hydroxymethylation data was available (unpublished) from entorhinal cortex DNA for 42 of the samples profiled in this study. DNA methylation and hydroxymethylation profiles were generated on the Illumina Infinium HumanMethylation450 BeadChip (Illumina Inc., CA, USA) (“Illumina 450K array”) using the TrueMethyl Array kit (Cambridge Epigenetix, Cambridge, UK). Profiles for both modifications were pre-processed, normalized and filtered according to a stringent standardised quality control pipeline, as described previously^50^ using the *WateRmelon*^68^ package in *R*. We identified probes on the array within 1kb of differentially acetylated peaks (FDR < 0.05) using the Bioconductor package *GenomicRanges*^62^. A total of 1,659 of the 4,162 FDR significant differentially acetylated peaks were located within 1kb of at least one CpG probe on the array, with a total of 6,838 probes mapping to the 1kb neighbourhood of these 1,649 peaks. For each CpG - peak pair we correlated the log fold change in H3K27ac between AD cases and controls to the difference in DNA methylation or hydroxymethylation in a linear model controlling for the same covariates as in the differential acetylation analysis. Moreover, we compared DNA methylation and hydroxymethylation between probes in vicinity of AD hyper - and hypoacetylated peaks, as well as those in vicinity of all background peaks and the whole microarray background using two sample t-tests.

## Accession codes

Raw data has been deposited in GEO under accession number GSE102538.

## Acknowledgements

This work was funded by US National Institutes of Health grant R01 AG036039 to J.M. S.J.M. and T.R. were funded by the EU-FP7 Marie Curie ITN EpiTrain (REA grant agreement no. 316758). Sequencing infrastructure was supported by a Wellcome Trust Multi User Equipment Award (WT101650MA) and Medical Research Council (MRC) Clinical Infrastructure Funding (MR/M008924/1). Generation of DNA hydroxymethylation data was funded by an Alzheimer’s Association US New Investigator Research Grant (grant number NIRG-14-320878) to K.L., and a grant from BRACE (Bristol Research into Alzheimer’s and Care of the Elderly) to K.L. The authors acknowledge the help of Dr Konrad Paszkiewicz at the University of Exeter Sequencing Service for advice on ChIP-seq experiments. We also acknowledge the help of Dr Vladimir Teif at the University of Essex in generating the UCSC Genome Browser tracks. Analysis was facilitated by access to the Genome high performance computing cluster at the University of Essex School of Biological Sciences.

## Author contributions

S.J.M., T.R. and K.M. conducted laboratory experiments. J.M., L.C.S and S.J.M. designed the study. J.M. supervised the project and obtained funding. S.J.M. undertook primary data analyses and bioinformatics, with analytical and computational input from L.C.S. and S.N. E.H. provided and pre-processed data for GWAS enrichment analyses. C.T. and S.A.-S. provided samples for analysis. K.L. and A.S. generated and pre-processed the DNA modification data. J.P. provided advice for the ChIP-seq analyses. S.J.M., J.M. and L.C.S. drafted the manuscript. All of the authors read and approved the final submission.

## References

1. Brookmeyer R, Johnson E, Ziegler-Graham K, Arrighi HM. Forecasting the global burden of Alzheimer’s disease. Alzheimers Dement 2007; 3:186–91.

2. Wenk GL. Neuropathologic changes in Alzheimer’s disease. J Clin Psychiatry 2003; 64 Suppl 9:7–10.

3. Hardy J, Selkoe DJ. The amyloid hypothesis of Alzheimer’s disease: progress and problems on the road to therapeutics. Science 2002; 297:353–6.

4. Karch CM, Cruchaga C, Goate AM. Alzheimer’s disease genetics: from the bench to the clinic. Neuron 2014; 83:11–26.

5. Reitz C, Brayne C, Mayeux R. Epidemiology of Alzheimer disease. Nat Rev Neurol 2011; 7:137–52.

6. Corder EH, Saunders AM, Strittmatter WJ, Schmechel DE, Gaskell PC, Small GW, Roses AD, Haines JL, Pericak-Vance MA. Gene dose of apolipoprotein E type 4 allele and the risk of Alzheimer’s disease in late onset families. Science 1993; 261:921–3.

7. Liu CC, Kanekiyo T, Xu H, Bu G. Apolipoprotein E and Alzheimer disease: risk, mechanisms and therapy. Nat Rev Neurol 2013; 9:106–18.

8. Lambert JC, Ibrahim-Verbaas CA, Harold D, Naj AC, Sims R, Bellenguez C, DeStafano AL, Bis JC, Beecham GW, Grenier-Boley B, et al. Meta-analysis of 74,046 individuals identifies 11 new susceptibility loci for Alzheimer’s disease. Nature genetics 2013; 45:1452

9. Lunnon K, Mill J. Epigenetic studies in Alzheimer’s disease: current findings, caveats, and considerations for future studies. Am J Med Genet B Neuropsychiatr Genet 2013; 162B:789–99.

10. Lunnon K, Smith R, Hannon E, De Jager PL, Srivastava G, Volta M, Troakes C, Al-Sarraj S, Burrage J, Macdonald R, et al. Methylomic profiling implicates cortical deregulation of ANK1 in Alzheimer’s disease. Nat Neurosci 2014; 17:1164–70.

11. De Jager PL, Srivastava G, Lunnon K, Burgess J, Schalkwyk LC, Yu L, Eaton ML, Keenan BT, Ernst J, McCabe C, et al. Alzheimer’s disease: early alterations in brain DNA methylation at ANK1, BIN1, RHBDF2 and other loci. Nat Neurosci 2014; 17:1156–63.

12. Creyghton MP, Cheng AW, Welstead GG, Kooistra T, Carey BW, Steine EJ, Hanna J, Lodato MA, Frampton GM, Sharp PA, et al. Histone H3K27ac separates active from poised enhancers and predicts developmental state. Proc Natl Acad Sci U S A 2010; 107:21931–6.

13. Kilgore M, Miller CA, Fass DM, Hennig KM, Haggarty SJ, Sweatt JD, Rumbaugh G. Inhibitors of class 1 histone deacetylases reverse contextual memory deficits in a mouse model of Alzheimer’s disease. Neuropsychopharmacology 2010; 35:870–80.

14. Cuadrado-Tejedor M, Garcia-Barroso C, Sanchez-Arias JA, Rabal O, Perez-Gonzalez M, Mederos S, Ugarte A, Franco R, Segura V, Perea G, et al. A First-in-Class Small-Molecule that Acts as a Dual Inhibitor of HDAC and PDE5 and that Rescues Hippocampal Synaptic Impairment in Alzheimer’s Disease Mice. Neuropsychopharmacology 2017; 42:524–39.

15. Fischer A. Targeting histone-modifications in Alzheimer’s disease. What is the evidence that this is a promising therapeutic avenue? Neuropharmacology 2014; 80:95–102.

16. Lu X, Wang L, Yu C, Yu D, Yu G. Histone Acetylation Modifiers in the Pathogenesis of Alzheimer’s Disease. Front Cell Neurosci 2015; 9:226.

17. Rao JS, Keleshian VL, Klein S, Rapoport SI. Epigenetic modifications in frontal cortex from Alzheimer’s disease and bipolar disorder patients. Transl Psychiatry 2012; 2:e132.

18. Zhang K, Schrag M, Crofton A, Trivedi R, Vinters H, Kirsch W. Targeted proteomics for quantification of histone acetylation in Alzheimer’s disease. Proteomics 2012; 12:1261–8.

19. Lu T, Aron L, Zullo J, Pan Y, Kim H, Chen Y, Yang TH, Kim HM, Drake D, Liu XS, et al. REST and stress resistance in ageing and Alzheimer’s disease. Nature 2014; 507:448–54.

20. Narayan PJ, Lill C, Faull R, Curtis MA, Dragunow M. Increased acetyl and total histone levels in post-mortem Alzheimer’s disease brain. Neurobiol Dis 2015; 74:281–94.

21. Sun W, Poschmann J, Cruz-Herrera Del Rosario R, Parikshak NN, Hajan HS, Kumar V, Ramasamy R, Belgard TG, Elanggovan B, Wong CC, et al. Histone Acetylome-wide Association Study of Autism Spectrum Disorder. Cell 2016; 167:1385–97 e11.

22. Kundaje A, Meuleman W, Ernst J, Bilenky M, Yen A, Heravi-Moussavi A, Kheradpour P, Zhang Z, Wang J, Ziller MJ, et al. Integrative analysis of 111 reference human epigenomes. Nature 2015; 518:317–30.

23. Anders S, Pyl PT, Huber W. HTSeq––a Python framework to work with high-throughput sequencing data. Bioinformatics 2015; 31:166–9.

24. Robinson MD, McCarthy DJ, Smyth GK. edgeR: a Bioconductor package for differential expression analysis of digital gene expression data. Bioinformatics 2010; 26:139-40.

25. Guintivano J, Aryee MJ, Kaminsky ZA. A cell epigenotype specific model for the correction of brain cellular heterogeneity bias and its application to age, brain region and major depression. Epigenetics : official journal of the DNA Methylation Society 2013; 8:290–302.

26. McLean CY, Bristor D, Hiller M, Clarke SL, Schaar BT, Lowe CB, Wenger AM, Bejerano G. GREAT improves functional interpretation of cis-regulatory regions. Nature biotechnology 2010; 28:495–501.

27. Thomas-Chollier M, Darbo E, Herrmann C, Defrance M, Thieffry D, van Helden J. A complete workflow for the analysis of full-size ChIP-seq (and similar) data sets using peak-motifs. Nat Protoc 2012; 7:1551–68.

28. Thomas-Chollier M, Herrmann C, Defrance M, Sand O, Thieffry D, van Helden J. RSAT peak-motifs: motif analysis in full-size ChIP-seq datasets. Nucleic Acids Res 2012; 40:e31.

29. Santpere G, Nieto M, Puig B, Ferrer I. Abnormal Sp1 transcription factor expression in Alzheimer disease and tauopathies. Neurosci Lett 2006; 397:30–4.

30. Citron BA, Dennis JS, Zeitlin RS, Echeverria V. Transcription factor Sp1 dysregulation in Alzheimer’s disease. J Neurosci Res 2008; 86:2499–504.

31. Ittner LM, Gotz J. Amyloid-beta and tau––a toxic pas de deux in Alzheimer’s disease. Nat Rev Neurosci 2011; 12:65–72.

32. Spillantini MG, Goedert M. Tau pathology and neurodegeneration. Lancet Neurol 2013; 12:609–22.

33. Ernst J, Kellis M. ChromHMM: automating chromatin-state discovery and characterization. Nat Methods 2012; 9:215–6.

34. Scheuner D, Eckman C, Jensen M, Song X, Citron M, Suzuki N, Bird TD, Hardy J, Hutton M, Kukull W, et al. Secreted amyloid beta-protein similar to that in the senile plaques of Alzheimer’s disease is increased in vivo by the presenilin 1 and 2 and APP mutations linked to familial Alzheimer’s disease. Nat Med 1996; 2:864–70.

35. Goate A, Chartier-Harlin MC, Mullan M, Brown J, Crawford F, Fidani L, Giuffra L, Haynes A, Irving N, James L, et al. Segregation of a missense mutation in the amyloid precursor protein gene with familial Alzheimer’s disease. Nature 1991; 349:704–6.

36. Cruchaga C, Haller G, Chakraverty S, Mayo K, Vallania FL, Mitra RD, Faber K, Williamson J, Bird T, Diaz-Arrastia R, et al. Rare variants in APP, PSEN1 and PSEN2 increase risk for AD in late-onset Alzheimer’s disease families. PloS one 2012; 7:e31039.

37. De Strooper B, Saftig P, Craessaerts K, Vanderstichele H, Guhde G, Annaert W, Von Figura K, Van Leuven F. Deficiency of presenilin-1 inhibits the normal cleavage of amyloid precursor protein. Nature 1998; 391:387–90.

38. Goate A, Hardy J. Twenty years of Alzheimer’s disease-causing mutations. J Neurochem 2012; 120 Suppl 1:3–8.

39. Crehan H, Holton P, Wray S, Pocock J, Guerreiro R, Hardy J. Complement receptor 1 (CR1) and Alzheimer’s disease. Immunobiology 2012; 217:244–50.

40. Heppner FL, Ransohoff RM, Becher B. Immune attack: the role of inflammation in Alzheimer disease. Nat Rev Neurosci 2015; 16:358–72.

41. Villegas-Llerena C, Phillips A, Garcia-Reitboeck P, Hardy J, Pocock JM. Microglial genes regulating neuroinflammation in the progression of Alzheimer’s disease. Curr Opin Neurobiol 2016; 36:74–81.

42. Ashburner M, Ball CA, Blake JA, Botstein D, Butler H, Cherry JM, Davis AP, Dolinski K, Dwight SS, Eppig JT, et al. Gene ontology: tool for the unification of biology. The Gene Ontology Consortium. Nature genetics 2000; 25:25–9.

43. Osborne JD, Flatow J, Holko M, Lin SM, Kibbe WA, Zhu LJ, Danila MI, Feng G, Chisholm RL. Annotating the human genome with Disease Ontology. BMC Genomics 2009; 10 Suppl 1:S6.

44. Robinson PN, Mundlos S. The human phenotype ontology. Clin Genet 2010; 77:525–34.

45. Jaeger S, Pietrzik CU. Functional role of lipoprotein receptors in Alzheimer’s disease. Curr Alzheimer Res 2008; 5:15–25.

46. Sun X, He G, Qing H, Zhou W, Dobie F, Cai F, Staufenbiel M, Huang LE, Song W. Hypoxia facilitates Alzheimer’s disease pathogenesis by up-regulating BACE1 gene expression. Proc Natl Acad Sci U S A 2006; 103:18727–32.

47. Zlokovic BV. Neurovascular pathways to neurodegeneration in Alzheimer’s disease and other disorders. Nat Rev Neurosci 2011; 12:723–38.

48. Warren JD, Rohrer JD, Rossor MN. Frontotemporal dementia. Bmj 2013; 347:f4827.

49. Limon A, Reyes-Ruiz JM, Miledi R. Loss of functional GABA(A) receptors in the Alzheimer diseased brain. Proc Natl Acad Sci U S A 2012; 109:10071–6.

50. Lunnon K, Hannon E, Smith RG, Dempster E, Wong C, Burrage J, Troakes C, Al-Sarraj S, Kepa A, Schalkwyk L, et al. Variation in 5-hydroxymethylcytosine across human cortex and cerebellum. Genome biology 2016; 17:27.

51. Kaas GA, Zhong C, Eason DE, Ross DL, Vachhani RV, Ming GL, King JR, Song H, Sweatt JD. TET1 controls CNS 5-methylcytosine hydroxylation, active DNA demethylation, gene transcription, and memory formation. Neuron 2013; 79:1086–93.

52. Kinde B, Gabel HW, Gilbert CS, Griffith EC, Greenberg ME. Reading the unique DNA methylation landscape of the brain: Non-CpG methylation, hydroxymethylation, and MeCP2. Proc Natl Acad Sci U S A 2015; 112:6800–6.

53. Selkoe DJ, Hardy J. The amyloid hypothesis of Alzheimer’s disease at 25 years. EMBO Mol Med 2016; 8:595–608.

54. Graff J, Tsai LH. Histone acetylation: molecular mnemonics on the chromatin. Nat Rev Neurosci 2013; 14:97–111.

55. Braak H, Braak E. Neuropathological stageing of Alzheimer-related changes. Acta Neuropathol 1991; 82:239–59.

56. Andrews S. FastQC: a quality control tool for high throughput sequence data.

57. Langmead B, Trapnell C, Pop M, Salzberg SL. Ultrafast and memory-efficient alignment of short DNA sequences to the human genome. Genome biology 2009; 10:R25.

58. Li H, Handsaker B, Wysoker A, Fennell T, Ruan J, Homer N, Marth G, Abecasis G, Durbin R, Genome Project Data Processing S. The Sequence Alignment/Map format and SAMtools. Bioinformatics 2009; 25:2078–9.

59. Quinlan AR, Hall IM. BEDTools: a flexible suite of utilities for comparing genomic features. Bioinformatics 2010; 26:841–2.

60. Zhang Y, Liu T, Meyer CA, Eeckhoute J, Johnson DS, Bernstein BE, Nusbaum C, Myers RM, Brown M, Li W, et al. Model-based analysis of ChIP-Seq (MACS). Genome biology 2008; 9:R137.

61. Love MI, Huber W, Anders S. Moderated estimation of fold change and dispersion for RNA-seq data with DESeq2. Genome biology 2014; 15:550.

62. Lawrence M, Huber W, Pages H, Aboyoun P, Carlson M, Gentleman R, Morgan MT, Carey VJ. Software for computing and annotating genomic ranges. PLoS Comput Biol 2013; 9:e1003118.

63. Lun AT, Chen Y, Smyth GK. It’s DE-licious: A Recipe for Differential Expression Analyses of RNA-seq Experiments Using Quasi-Likelihood Methods in edgeR. Methods Mol Biol 2016; 1418:391–416.

64. Mathelier A, Fornes O, Arenillas DJ, Chen CY, Denay G, Lee J, Shi W, Shyr C, Tan G, Worsley-Hunt R, et al. JASPAR 2016: a major expansion and update of the open-access database of transcription factor binding profiles. Nucleic Acids Res 2016; 44:D110–5.

65. Heinz S, Benner C, Spann N, Bertolino E, Lin YC, Laslo P, Cheng JX, Murre C, Singh H, Glass CK. Simple combinations of lineage-determining transcription factors prime cisregulatory elements required for macrophage and B cell identities. Mol Cell 2010; 38:576-89.

66. Chang CC, Chow CC, Tellier LC, Vattikuti S, Purcell SM, Lee JJ. Second-generation PLINK: rising to the challenge of larger and richer datasets. Gigascience 2015; 4:7.

67. Hannon E, Dempster E, Viana J, Burrage J, Smith AR, Macdonald R, St Clair D, Mustard C, Breen G, Therman S, et al. An integrated genetic-epigenetic analysis of schizophrenia: evidence for co-localization of genetic associations and differential DNA methylation. Genome biology 2016; 17:176.

68. Pidsley R, CC YW, Volta M, Lunnon K, Mill J, Schalkwyk LC. A data-driven approach to preprocessing Illumina 450K methylation array data. BMC Genomics 2013; 14:293.

